# Phase transitions in tumor growth: II prostate cancer cell lines

**DOI:** 10.1101/011189

**Authors:** J.A. Llanos-Pérez, A. Betancourt-Mar, M. P. De Miguel, E. Izquierdo-Kulich, M. Royuela-García, E. Tejera, J.M. Nieto-Villar

## Abstract

We propose a mechanism for prostate cancer cell lines growth, e.g., LNCaP and PC3 based on a Gompertz dynamics. This growth exhibits a multifractal behavior and a “second order” phase transition. Finally, it was found that the cellular line PC3 exhibits a higher value the entropy production rate compared to LNCaP, which is indicative of the robustness of PC3, over to LNCaP and may be a quantitative index of metastatic potential tumors.

**Highlights:**

- Cancer is an open, complex, dynamic and self-organizing system.
- Prostate cancer cell lines growth follows a Gompertz dynamics
- Prostate cancer cell lines exhibit a multifractal behavior
- The entropy production rate may be considered as metastatic potential marker

## Introduction

Cancer is a generic name given to a group of malignant cells which have lost their specialization and control over normal growth. This group of malignant cells could be considered as a nonlinear dynamical system, self-organized in time and space, far from thermodynamic equilibrium, exhibiting high complexity [1], robustness [2] and adaptability [3].

Transitions phenomena far from thermodynamic equilibrium are a consequence of bifurcations and correlations which may have relevance it’s the macroscopic behavior of the tumor. On the other hand, bifurcations in dynamical systems have an analogous function to phase transitions in the vicinity of equilibrium. These result from on increment and amplification at the macroscopic levels of microscopic fluctuations. It is the main mechanism of self-organization, and, consequently, of complexity at the macroscopic level [4].

Recently, we had shown [5] that the transition from healthy cells to malignant cells during to avascular growth can be described by a second-order phase transition through either logistic or Gompertz dynamic equations equivalently.

The goal of this work is to extend the thermodynamics formalism previously developed [5] to the dynamical behavior of the prostate tumor cell lines, such as, a example LNCaP and PC3.

The manuscript is organized as follow: Section 2 describes the experimental part: the growth dynamics of prostate tumor cell lines LNCaP and PC3 and the dynamical behavior of the fractal dimension of the cell lines patterns. In section 3 we develop a thermodynamic framework, based on the entropy production rate. Finally, some concluding remarks are presented.

## 2. Experimental Part: A dynamic behavior and the fractal dimension of the prostate tumor cell lines, LNCaP and PC3

The prostate tumor cell lines, serve as models of tumor growth [6] as well as for the development of various therapies [7]. Human prostatic cancer cells LNCaP (CRL-1740) and PC3 (CRL-1435), supplied by American Type Culture Collection (ATCC, Rockville, MD), were routinely cultured [8]. Images were taken over an 8 day period; all images have a magnification of 40X.

The system in question is a 2*D* region with characteristic length *L* in which initially there are very few tumor cells. The cell density increases with time due to the proliferation of these (Figure 1).

**Fig. 1.**
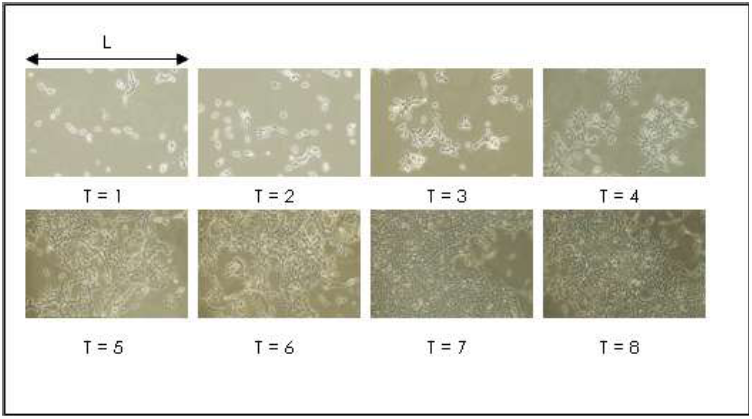
*in vitro* growth of prostate tumor cell lines. *T* is time in days.

The morphology observed in this region has fractal nature as a result of the stochastic nature of the mitosis and apoptosis processes that occur at the level of single cells [9]. Thus we have

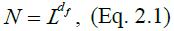

where *N* is the number of total cells in the region of characteristic size *L* and *d*_*f*_ fractal dimension of the contour. The Euleriana of (Eq. 2.1) is given by

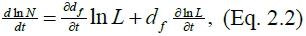

In previous work [5] we show that the temporal variation of the total number of cells *N* can be described equivalently through a logistic or Gompertz dynamical equation. If the dynamical system exhibits temporal multifractality the first term on the right of equality (2.2) is different from zero. In this way we have 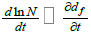 whereby the fractal dimension variation in time can be described, in principle, by expression equivalent to the Gompertz equation, such that

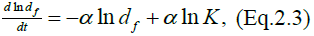

where, 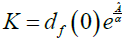 is the carrying capacity and *Â* and *α* are empirical constants in the Gompertz equation [10]. Equation (2.3) has the analytical solution

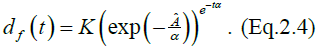

Equation (2.4), for a dynamical system that exhibits temporal multifractal behavior, provides the functional dependence of *d*_*f*_ with the time. In Fig. 2, the time dependence of the fractal dimension is shown *d*_*f*_ for the dynamical behavior of the prostate tumor cell lines, LNCaP and PC3 respectively. The morphological characterization of each image was obtained by the software ImageJ 1.43 [11].

**Fig. 2.**
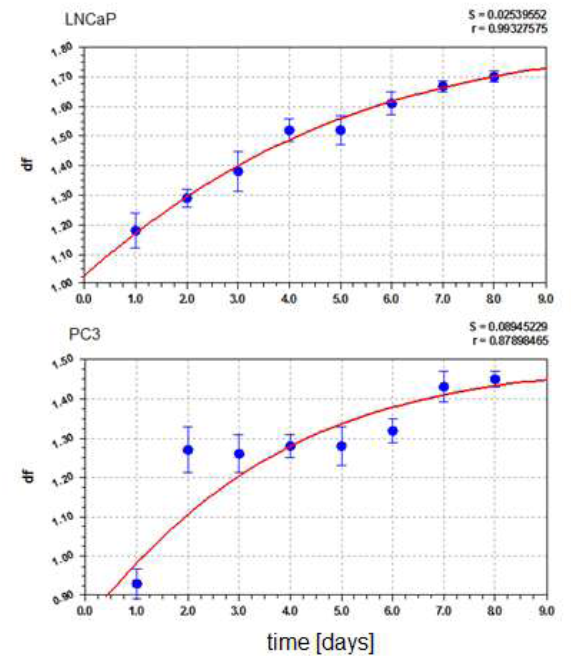
Dependence of fractal dimension *d*_*f*_ with the growth time of the prostate tumor cell lines: LNCaP (*Â* = 0.2524, *α* = 0.1466, *K* = 1.8384) and PC3 (*Â* = 0.3335, *α* = 0.1944, *K* = 1.4924).

These graphs exhibit a multifractal behavior, in distinction to other cell lines showing fractal behavior [12]. A satisfactory dynamic fit is achieved through the formula (1.4).

As shown in a previous work [9], the fractal dimension *d _f_* can be given as a function of the quotient between mitosis *V_m_* and apoptosis *V_a_* rates [13], which quantify the tumor capacity to invade and infiltrate healthy tissue:

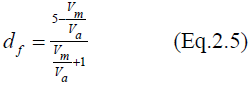

Substituting the temporal dependence of *d*_*f*_ for each of the cells (equation 2.4), in the formula (2.5) we obtain the time dependence of the 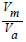 ratio as illustrated in Figure 3:

**Fig. 3.**
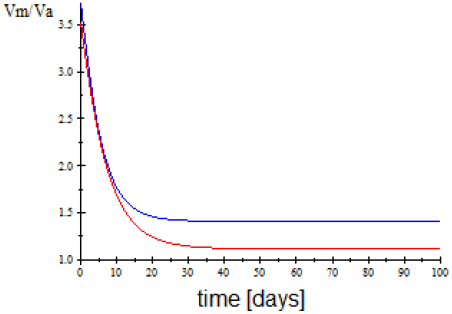
time dependence of the quotient between mitosis *V*_*m*_ and apoptosis *V*_*a*_ rates: PC3 (blue), LnCap (red).

As shown in fig.3, the cell line PC3, which is known to have an increased invasive ability and is more aggressive as compared to the LnCap [14,15], shows a higher value quotient between mitosis *V*_*m*_ and apoptosis *V*_*a*_ rates.

## 3. Thermodynamics approach

In recent years various authors [16, 17] have argued that through a thermodynamic approach to cancer as nonlinear dynamical system that self-organizes out of equilibrium we can not only understand its dynamics and complexity but also its robustness.

We have recently demonstrated that the entropy production rate 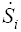 for avascular tumor growth is related with the cancer robustness [5,18] and can be expressed as a function of fractal dimension *d*_*f*_ as:

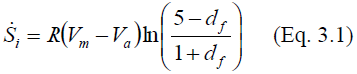

where *R* is the gas constant. In formula (3.1), two properties related to on tumor growth are included: the first is their growth rate, which is associated with their invasiveness; the second is its complexity, a morphology characteristic as the fractal dimension of the tumor interface, which quantifies the tumor capacity to invade and infiltrate healthy tissue.

Considering Eq. (2.4) and Eq. (2.5), and the temporal dependence of the fractal dimension *d*_*f*_ of the dynamical behavior of the prostate tumor cell lines, LNCaP and PC3 (Fig. 2), and substituting in Eq. (3.1) the functional dependence the entropy production rate 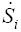 is obtained in the time, as shown in Fig. 4:

**Fig 4.**
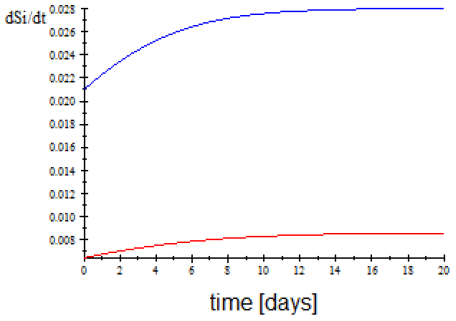
The entropy production rate 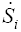 for the different the prostate tumor cell lines: PC3 (blue), LnCap (red).

As shown in Fig.4, cell line PC3 shows a greater value far the entropy production rate compared to a LnCap. These results corroborate, from the new point of thermodynamics, what others studies have shown far the cell line PC3, that they are more malignant, aggressive, have a higher metastatic potential and are more resistant to treatment compared to LNCap [19].

## Conclusion

In summary, in this paper we found:

1. We propose a mechanism for prostate cancer cell lines growth, LNCaP and PC3, based on a Gompertz dynamics.
2. The growth of the cell lines, LNCaP and PC3, exhibit a multifractal behavior and the “second order” phase transition.
3. It was found that the cellular line PC3 exhibits a greater value of the entropy production rate compared to LNCaP, which is indicative of the robustness of PC3 in comparison to LNCaP and this may be used a quantitative index of the metastatic potential of tumors.

The current theoretical framework for prostate cancer cell lines growth, LNCaP and PC3, will hopefully provide a better understanding of cancer and contribute to improvements in cancer treatment.

## Acknowledgements

We would like to thank Prof. Jacques Rieumont for support and encouragement of this research. Dr. K. Michaelian Michelian from Instituto de Física de La UNAM México for comments and interesting suggestions. This work was supported by a grant, in part by the project SAF2010-19230 of the Ministerio Español de Ciencia e Innovación al Laboratorio de Ingeniería Celular-IdiPAZ, as well as Biomedicine and Biotechnology, University of Alcalá; and Mexican Institute of Complex Systems, Geo-estratos S.A. de México.

